# Gene recoding by synonymous mutations creates promiscuous intragenic transcription initiation in mycobacteria

**DOI:** 10.1101/2023.03.17.532606

**Authors:** Nuri K. Hegelmeyer, Mary L. Previti, Joshua Andrade, Raditya Utama, Richard J. Sejour, Justin Gardin, Stephanie Muller, Steven Ketchum, Alisa Yurovsky, Bruce Futcher, Sara Goodwin, Beatrix Ueberheide, Jessica C. Seeliger

**Author notes:** Address correspondence to Jessica C. Seeliger. Josh Andrade (6sense, New York, New York), Justin Gardin (Gingko Bioworks, Inc., Boston, Massachusetts).

## Abstract

Each genome encodes some codons more frequently than their synonyms (codon usage bias), but codons are also arranged more frequently into specific pairs (codon pair bias). Recoding viral genomes and yeast or bacterial genes with non-optimal codon pairs has been shown to decrease gene expression. Gene expression is thus importantly regulated not only by the use of particular codons but by their proper juxtaposition. We therefore hypothesized that non-optimal codon pairing could likewise attenuate *Mtb* genes. We explored the role of codon pair bias by recoding *Mtb* genes (*rpoB, mmpL3, ndh*) and assessing their expression in the closely related and tractable model organism *M. smegmatis*. To our surprise, recoding caused the expression of multiple smaller protein isoforms from all three genes. We confirmed that these smaller proteins were not due to protein degradation, but instead issued from new transcription initiation sites positioned within the open reading frame. New transcripts gave rise to intragenic translation initiation sites, which in turn led to the expression of smaller proteins. We next identified the nucleotide changes associated with these new sites of transcription and translation. Our results demonstrated that apparently benign, synonymous changes can drastically alter gene expression in mycobacteria. More generally, our work expands our understanding of the codon-level parameters that control translation and transcription initiation.

**IMPORTANCE:** *Mycobacterium tuberculosis* (*Mtb*) is the causative agent of tuberculosis, one of the deadliest infectious diseases worldwide. Previous studies have established that synonymous recoding to introduce rare codon pairings can attenuate viral pathogens. We hypothesized that non-optimal codon pairing could be an effective strategy for attenuating gene expression to create a live vaccine for *Mtb*. We instead discovered that these synonymous changes enabled the transcription of functional mRNA that initiated in the middle of the open reading frame and from which many smaller protein products were expressed. To our knowledge, this is the first report that synonymous recoding of a gene in any organism can create or induce intragenic transcription start sites.

## INTRODUCTION

Across the tree of life, genomes encode some codons more commonly than others, a preference known as codon usage bias. Independently of this well-known phenomenon, each organism also exhibits preference for certain codon pairs. First termed twenty-five years ago, codon *pair* bias (codon pairB) describes the preferential use of some hexameric codon pairs over other synonymous pairs (1). Not long after the first genomes were sequenced, early bioinformatics studies reported on the prevalence of codon pairB, which they found in diverse coding genomes, from eukaryotes to bacteria (2–4).

As the more extensively studied phenomenon, codon usage bias has been applied broadly across organisms towards optimizing or, less often, attenuating gene expression (5). In contrast, most codon pairB studies to date have sought to impair gene expression, and predominantly in viral pathogens. Prior codon pairB studies have shown that gene expression can be attenuated by synonymously recoding viral genomes so they include more rare codon pairs without perturbing codon usage, in a process called codon pair deoptimization. One of the first reports on the functional consequences of codon pairB demonstrated that synonymously recoding the poliovirus genome by codon pair deoptimization suppressed viral gene expression, reduced viral replication, and yielded a virus that protected mice against polio when used as a vaccine (6). Subsequent studies investigating gene attenuation by altering codon pairs suggest that the mechanism of attenuation is most likely multifactorial. In yeast, deoptimizing some codon pairs caused changes in translation efficiency (7,8), while recoding an influenza gene decreased the half-life of the viral mRNA (9).Though the exact mechanism remains elusive, synonymous recoding by codon pair deoptimization has been applied to attenuate several other pathogenic viruses, including influenza, Zika virus, and the recently emergent coronavirus SARS-CoV-2 (10–18).

In contrast, there has been only one study on codon pair deoptimization to attenuate a bacterial pathogen. Coleman *et al*. sought to attenuate *Streptococcus pneumoniae* by synonymously recoding a single gene, the virulence factor pneumolysin (*ply*) (19). Recoding *ply* led to reduced Ply expression and rendered the mutant less virulent than the wild type. However, the mechanism by which deoptimizing codon pair bias compromises bacterial gene expression has not, to our knowledge, been explored.

*Mycobacterium tuberculosis* (*Mtb)* is the causative pathogen of the ancient and deadly infectious disease tuberculosis (TB). TB caused an estimated 1.3 million deaths in 2021 and remains among the leading infectious causes of death worldwide, second only to COVID-19 (20). The attenuated strain *Mycobacterium bovis bacille* Calmette-Guérin, or BCG, is given worldwide to newborns as a vaccine against severe TB, but is not considered protective against pulmonary TB, which is the primary cause of mortality. Based on previous studies with viral pathogens and *S. pneumoniae*, we hypothesized that synonymously recoding essential genes in *Mtb* would attenuate protein expression and thereby reduce bacterial survival and pathogenicity. We chose to recode the codon pair bias in several *Mtb* genes: the RNA polymerase β-subunit *rpoB* (*rv0667*) and the trehalose monomycolate exporter *mmpL3* (*rv0206c*), which are essential in normoxic conditions (21–23), and the type-II NADH dehydrogenase *ndh* (*rv1854c*), which is essential under hypoxia and on host-related carbon sources (22,24). We first sought to confirm that recoding attenuated protein expression but found instead that recoding results in the expression of many smaller protein isoforms that appeared to initiate from recoded portions of the open reading frame (ORF). This unexpected result prompted us to investigate mechanisms of pervasive intragenic expression.

Here, we report the impact of synonymous recoding on translation and transcription initiation in *Mtb* genes. We examined the expression of wildtype and recoded *Mtb* genes and found that synonymous mutations can give rise to intragenic transcription start sites (TSS) that produce functional mRNA. We further demonstrated that the smaller protein isoforms expressed from transcripts initiating from *de novo* intragenic promoters. Finally, we uncovered evidence that the coding genome exhibits bias against codon pair or nucleotide hexamer biases that can cause transcription, indicating that these higher order patterns in the genetic code may serve to moderate gene expression. Taken together, our findings reveal that certain synonymous changes can drastically alter the way genes are expressed in mycobacteria, further supporting the growing consensus that synonymous mutations are not always silent.

## RESULTS

### Synonymous mutations to introduce rare codon pairs cause expression of additional protein products

Previous work on codon pair bias and gene expression in viruses and *S. pneumoniae* (6,19) led us to hypothesize that synonymous recoding to introduce rare pairs of codons, also known as codon pair deoptimization, would similarly attenuate gene expression in mycobacteria. We further hypothesized that recoding longer gene segments would lead to a higher degree of attenuation.

The codon pair bias of a sequence is described using codon pair scores (**Supplemental Figure S1A**). A codon pair score is derived from the natural log transformation of the ratio of the observed frequency of a given codon pair (*i.e.*, in-frame hexamer) to the expected frequency of a given codon pair, controlling for the number of times the corresponding dipeptide is encoded across the entire coding genome (**Supplemental Figure S1B**). Positive scores indicate “enriched” codon pairs, relative to other codon pairs encoding the same dipeptide, while negative scores describe “depleted” pairs. For example, the hexamer TTA-CGT represents the most enriched codon pair for the Leu-Arg dipeptide in *Mtb* and has a codon pair score of 0.724, whereas the least common Leu-Arg dipeptide in *Mtb* is CTC-CGG with a codon pair score of −1.430. A gene can then be described by a codon pair bias score, which is the mean of codon pair scores for all in-frame hexamers (**Supplemental Figure S1C**, see also: Methods). Substituting CTC-CGG for TTA-CGT does not change the amino acid sequence—and is thus a synonymous mutation—but does replace a frequently occurring codon pair with a rarer one and thereby decreases the codon pair bias score for a gene. Thus, codon pair bias score is commonly used to indicate how far codon pair bias within a single coding sequence deviates from the coding genome as a whole; however, we note since codon pair bias score is an average, a gene can contain very rare codon pairs whose effects on the score are offset by frequently occurring codon pairs elsewhere in the sequence. Using codon pair bias score as a metric thus assumes that the effects of codon pair bias on gene expression are aggregate and not individual. As previously mentioned, synonymously rearranging the codons within a gene to increase the number of rare codon pairs is a process known as codon pair deoptimization. Hereafter, we refer to this process as “recoding.”

In a pilot study we selected several genes that are required for the survival of *Mtb* under normoxic (RNA polymerase β-subunit *rpoB*; trehalose monomycolate transporter *mmpL3*) or hypoxic (type-II NADH dehydrogenase *ndh*) culture conditions. For this test system we recoded the genes by shuffling codons within each reading frame to maximize the occurrence of rare codon pairs without significantly altering codon usage. For each gene, we further generated up to three variants in which we recoded different length segments of the ORF starting from the 3′ end (**Figure 1A**). Each variant was independently recoded and so represents a unique sequence, and the codon pair bias score (*i.e.*, the mean codon pair score of each gene) was deoptimized relative to bias in *Mtb* coding genome (see Methods). For consistent expression, each variant encoded a C-terminal 3XFLAG epitope fusion and was expressed from a multicopy episomal plasmid in which transcription was driven by the constitutive *groEL* promoter (P*groEL*) (25). For initial experiments assessing protein expression, these constructs were transformed into the fast-growing, non-pathogenic species *M. smegmatis* (*Msm*).

**Figure 1.**
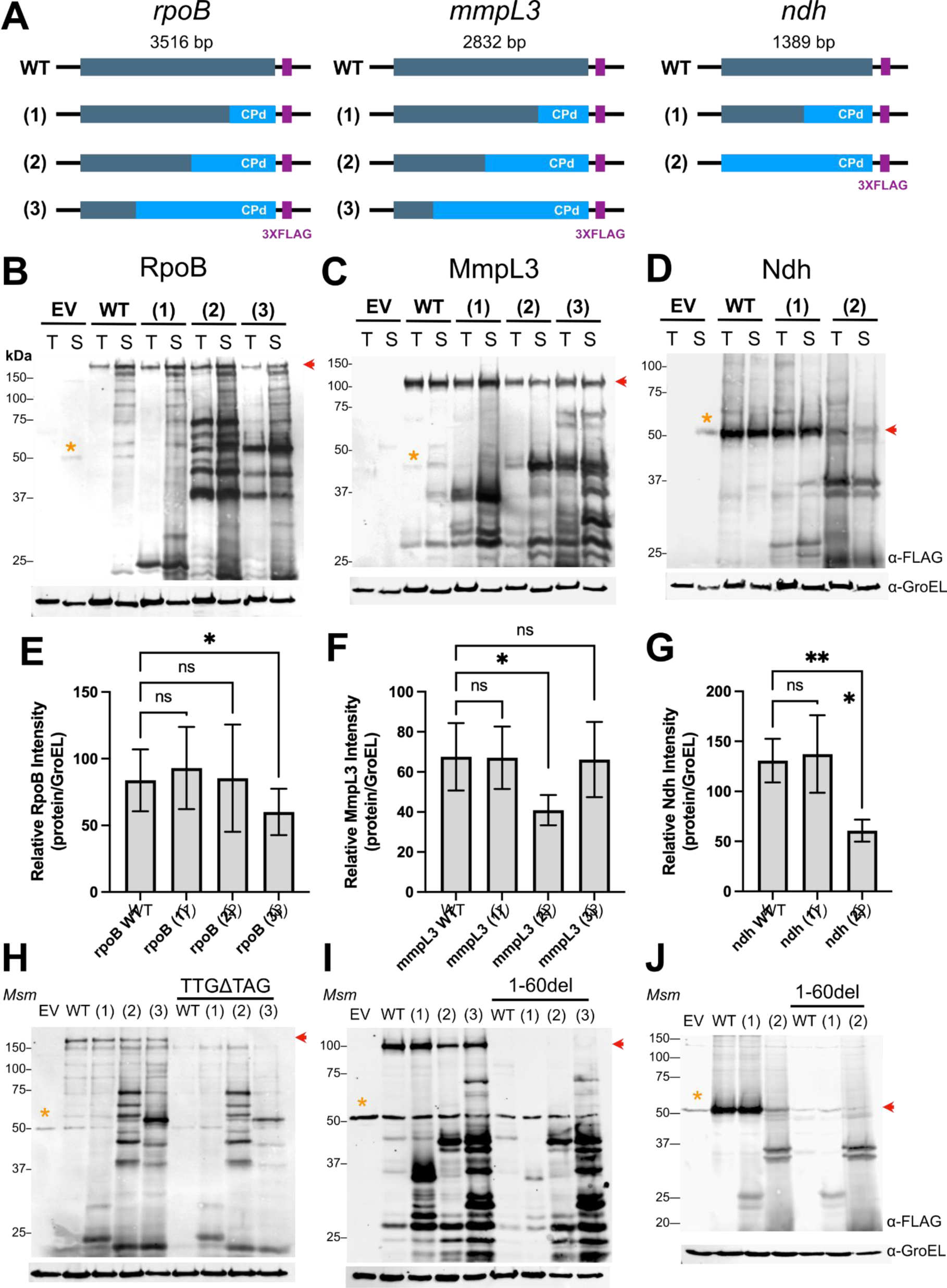
Synonymous mutation by codon pair deoptimization causes expression of smaller peptides in mycobacteria. (A) Diagram of the strategy to codon-pair deoptimize *Mtb* genes and assess their expression. Different lengths of *rpoB*, *mmpL3*, and *ndh* sequences were recoded starting from the 3′ end and fused with a 3XFLAG peptide epitope at the C-terminus. Anti-FLAG immunoblots of total lysates from *Mtb* (T) or *Msm* (S) carrying wild-type and recoded *Mtb* genes (B) *rpoB*, (C) *mmpL3*, or (D) *ndh* expressed from a multi-copy episomal plasmid. (E, F, G) Quantification of full-length protein from each *Msm* strain in (B, C, D) normalized to a GroEL loading control. Data represent the average ± S.D. of n=5 independent experiments. Repeat Measures One-Way ANOVA with Dunnett multiple comparison analysis was performed to test significance. * p>0.05, *** p>0.002. Anti-FLAG immunoblots of *Msm* strains in which (H) the start codon of *rpoB* was replaced with a stop codon (TTGΔTAG) or (I,J) the first 60 nt were deleted from *mmpL3* and *ndh*. Immunoblots are representative of ≥3 independent experiments. Red arrows indicate the expected migration of the full-length proteins. Yellow asterisks indicate a non-specific FLAG cross-reacting band in *Msm* total lysates.

Contrary to our hypotheses, variants encoding segments with more rare codon pairs did not consistently lead to decreased protein expression (**Figure 1B-G**). Instead, a wholly unexpected and striking feature was the detection of multiple peptides at molecular weights lower than that of the corresponding full-length protein (**Figure 1B-D**). This phenomenon appeared to be general, as the additional protein products were observed for all recoded variants of all three genes. Also, the additional proteins were not an artifact of expression of *Mtb* genes (i) in the heterologous host *Msm*, (ii) from multiple gene copies, or (iii) under selection with the translational inhibitor hygromycin, as the pattern and relative intensities of the additional proteins were similar when the variants were expressed in *Mtb* or *Msm* (**Figure 1B-D**) or from a single stably integrated gene copy and in the absence of hygromycin (**Supplemental Figure S2**). Importantly, these results supported our further use of the experimentally tractable, fast-growing, and non-pathogenic species *Msm* to study the effects of recoding *Mtb* genes. Lastly, the additional proteins appeared to derive directly from the recoded gene segment, as for each variant, the size range of the new products corresponded qualitatively with the length of the recoded segment.

### Smaller proteins express from new translation initiation sites within recoded gene regions

We noted that the additional proteins must be in-frame isoforms since they were detected by anti-FLAG immunoblot and therefore must contain a C-terminal FLAG tag. Further, they are unlikely the product of degradation, since every gene variant encodes the wild-type amino acid sequence and there is no known mechanism by which a protease can discriminate between identical proteins. Nevertheless, to confirm that the smaller isoforms were not degradation products, we pursued two strategies to preclude translation of the full-length protein. First, we substituted the predicted start codon with a stop codon (TTGΔTAG). This mutation abolished expression of full-length RpoB, but not the smaller isoforms (**Figure 1H**). For Ndh and MmpL3, replacing the start codon with the TAG stop codon did not abolish expression of the full-length Ndh or MmpL3 (**Supplemental Figure S3**). We speculated that apparent full-length protein was still expressed because the actual (or additional) translation initial site(s) (TIS) were located just downstream of the predicted start codon. Thus, for Ndh and MmpL3 we created additional variants in which we deleted the first 60 nucleotides (20 aa) of the ORF, including the start codon (Δ1-60nt). This deletion abolished expression of full-length Ndh and MmpL3, but not that of the corresponding smaller isoforms (**Figure 2I, J**). Overall, these results confirmed that smaller proteins were not the result of post-translational cleavage. We therefore reasoned that they must originate from TIS within the ORF and, more specifically, from within the recoded segments.

**Figure 2.**
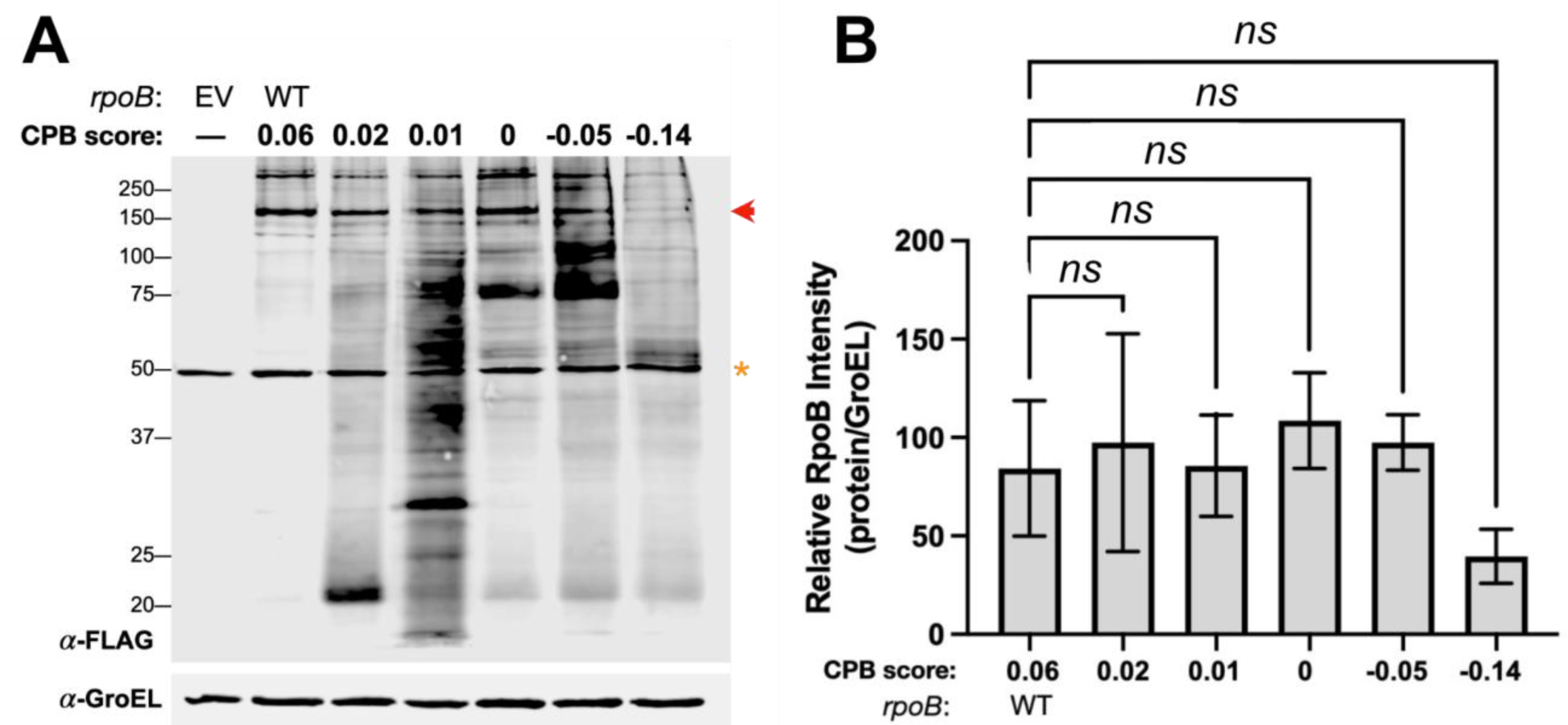
The number of smaller proteins and the expression of full-length protein varies but does not correlate with codon pair bias score. (A) Anti-FLAG immunoblot of *Msm* strains carrying wild-type *Mtb rpoB* or recoded variants with the indicated codon pairB scores. Red arrow indicates the expected migration of full-length RpoB. Yellow asterisk indicates a non-specific FLAG cross-reacting band in *Msm* total lysates. Immunoblot is representative of 3 independent experiments. (B) Quantification of full-length RpoB normalized to a GroEL loading control. Repeat Measures One-Way ANOVA with Dunnett multiple comparison analysis was performed to determine significance in n=3 independent experiments. All pairwise comparisons to WT were not significant (ns; p>0.05).

### Codon pair bias score of gene does not determine size or number of smaller proteins expressed

We observed no clear correlation between the length of gene recoded and quantity of smaller proteins expressed as represented by unique bands on immunoblot (**Figure 1B-D**). However, the question remained: If the recoded length is controlled, does the codon pair bias score of a recoded gene influence the sizes and quantity of smaller proteins produced? In other words, does intragenic expression arise from global changes across the gene or to local instances of rare codon pairs? To answer this question, we generated additional codon pair bias score variants, this time recoding the same sequence segment as in *rpoB*(3) (2394 of 3516 bp), which had the longest recoded segment and thus the highest theoretical probability of achieving a wide range of codon pair bias scores upon recoding. Starting from the wild-type sequence, we recoded this segment, this time by generating 100,000 randomly shuffled sequences. From this pool we selected sequences with a range of codon pair bias scores between those of *rpoB*(3) (−0.14) and the wild type (0.06).

We then assessed protein expression from these sequences to determine whether the levels of smaller protein isoforms could be tuned by codon pair bias score. As was the case when different sequence lengths were recoded, no obvious correlation was observed between the codon pair bias score and expression of full-length RpoB (**Figure 3**). Furthermore, no clear correlation was observed between codon pair bias score and the number, size, and amounts of additional protein products. These results suggested that the codon pair bias score, which describes the cumulative effect of codon shuffling, does not account for the production of smaller proteins, which we then postulated arises from local and particular sequence changes.

**Figure 3.**
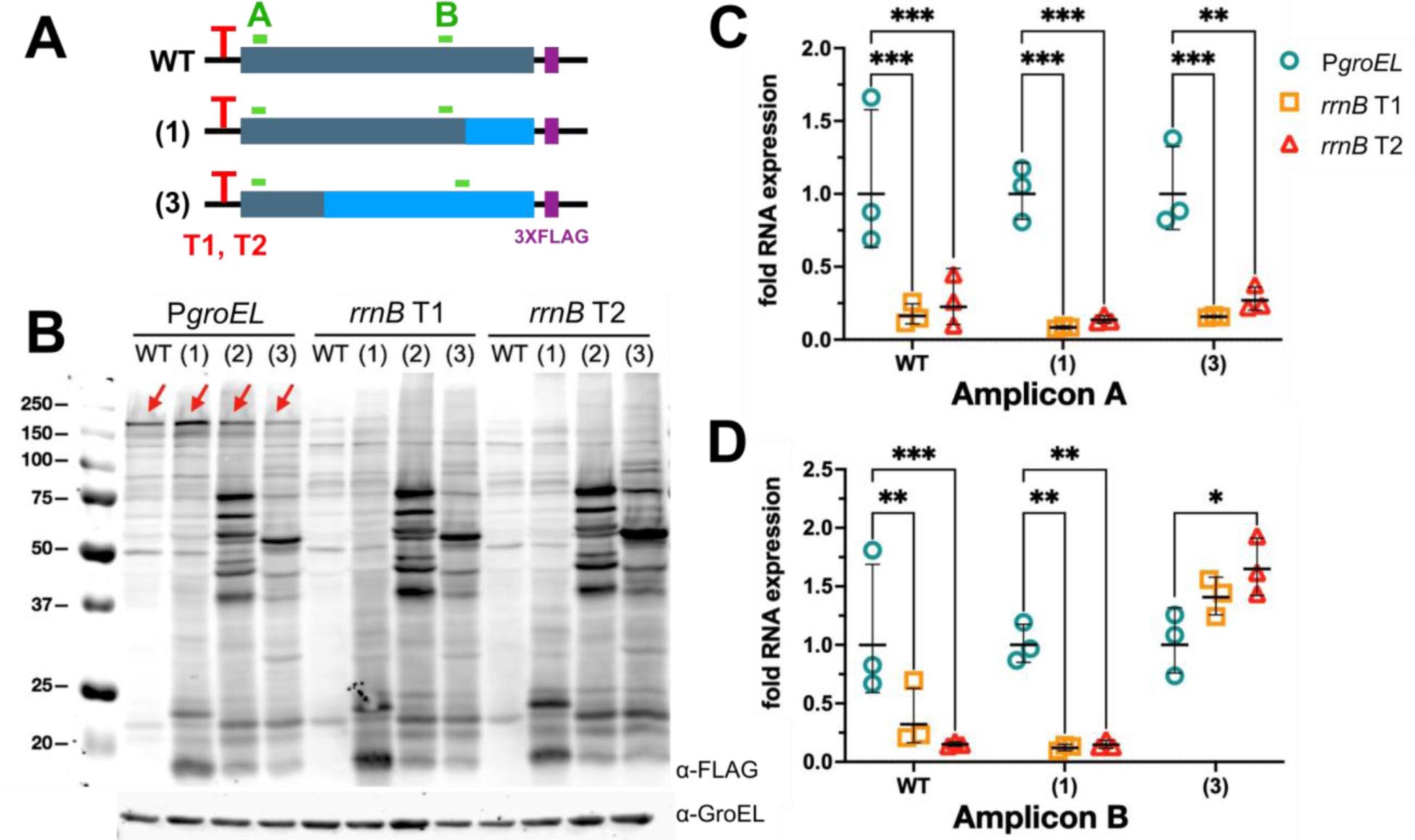
Recoded gene variants express transcripts that initiate from within recoded regions. (A) Diagram of *rpoB* genes in which the *groEL* promoter (P*groEL*) was substituted by the *E. coli rrnB* T1 or T2 terminator loop (red “T”) to repress full-length transcript. Amplicon A probes a 5′ segment encoding the wild-type sequence in all variants. Amplicon B probes a 3′ segment that is wild-type in WT and (1) but recoded in (3). (B) Anti-FLAG immunoblot *Msm* strains carrying WT *rpoB* or recoded variants. GroEL was used as a loading control. Immunoblot is represenstative of n=3 independent experiments. (C, D) qPCR of amplicons A and B for *Msm* strains carrying WT *rpoB* or recoded variants. Data and average ± S.D. from n=3 independent experiments are shown. Two-way ANOVA was performed to determine significant (* p<0.05; ** p<0.02; *** p<0.002).

### Synonymous recoding causes the transcription of mRNA that initiate in the middle of the open reading frame and are translated into smaller proteins

Having confirmed that recoding introduces additional TIS in the middle of the ORF, we next asked whether changes in transcription contribute to these new sites of protein synthesis. We hypothesized that introducing rare codon pairs can result in new transcriptional start sites (TSS). These new, shorter mRNAs can contain new translational initiation sites and lead to the expression of smaller proteins. To test this hypothesis, we sought to prevent production of the full-length transcript by replacing the *groEL* promoter with the *E. coli rrnB* T1 or T2 terminator loops, which function as strong transcriptional terminators in *M. smegmatis* (**Figure 3A**) (26). Immunoblot analysis showed that, as expected, expression of the full-length RpoB was reduced or abolished, but smaller isoforms persisted (**Figure 3B**). We confirmed that either terminator T1 or T2 strongly inhibited transcription of full-length *rpoB* mRNA using quantitative PCR of a 5′ amplicon common to all variants and the wild type (**Figure 3C**). In contrast, qPCR of a 3′ amplicon revealed that the recoded region of *rpoB*(3) was still transcribed (**Figure 3D**). We thus concluded that recoding using synonymous mutations introduced TSS within the ORF.

### 5′-end sequencing confirms transcription initiation within recoded regions

What specific sequence changes within recoded regions give rise to intragenic TSS and TIS? To answer this question we mapped the 5′ ends of mRNA transcripts by 5′ rapid amplification of cDNA ends (RACE), a method that selectively amplifies TSS. We hypothesized that RACE would identify more internal TSS (iTSS) in the recoded genes than in the wild type, as this would corroborate our qPCR data indicating that recoding creates new functional intragenic starts that give rise to smaller protein isoforms. First, we detected iTSS arising from the wild-type gene —a plausible finding since 5′-end mapping transcriptomics have shown that iTSS occur endogenously in mycobacteria (27–29). In addition and in support of our hypothesis we found that there were more iTSS in recoded variants—and specifically in the recoded portions of variant genes (**Figure 4**). The number of iTSS appeared to correspond with the length of recoded sequence such that more intragenic starts were mapped to *rpoB*(3) *vs. rpoB*(2). However, iTSS in recoded *rpoB*(3) were farther apart than in *rpoB*(2), consistent with immunoblots showing more smaller proteins expressed from the latter (**Figure 1B**).

**Figure 4.**
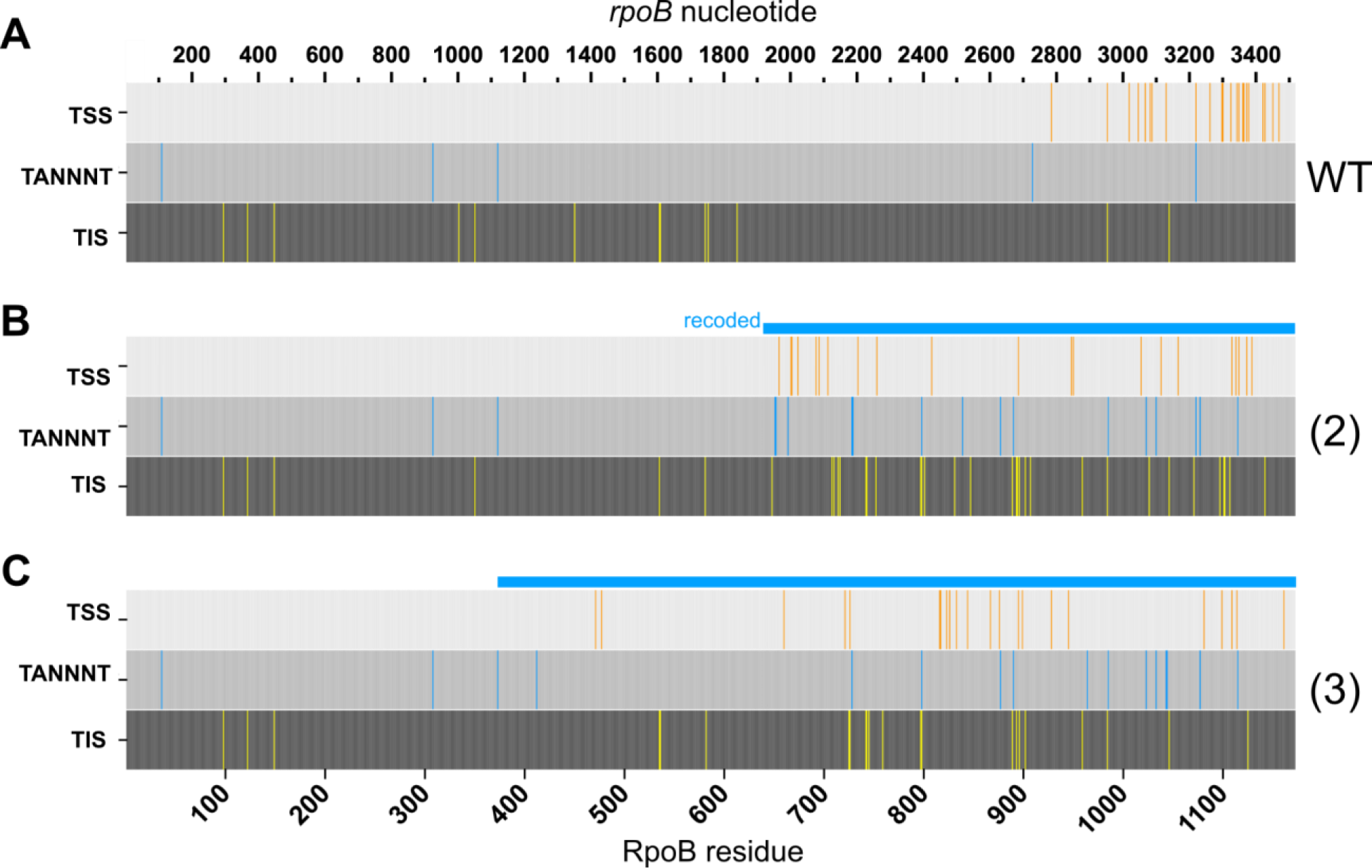
Gene recoding gives rise to a higher incidence of intragenic transcription and translation initiation. Transcription start sites (TSS) identified by 5′ RACE and translation initiation sites (TIS) identified by detection of N-terminally modified peptides for *Msm* strains carrying (A) wild-type *rpoB,* (B) variant (2), or (C) variant (3). The positions of TANNNT transcription initiation motifs are also indicated. Nucleotide position is indicated on the top axis; amino acid position on the bottom axis. The light blue bar above each gene indicates the recoded region. TSS were detected in at least 2 of 3 independent experiments. TIS were detected in 2 independent experiments.

In summary, we found that portions of the gene containing more rare codon pairs gave rise to more iTSS. This result supports a secondary hypothesis that codon pair bias reflects the depletion of codon pairs that stimulate transcription initiation.

### codon pair deoptimization increases the number of transcription motifs appearing intragenically

Mycobacterial TSS are typically preceded by known enhancers (*i.e.*, TANNNT) and begin with a purine (27–29). If introducing new motifs such at TANNNT or other enhancer motifs is in fact how codon pair deoptimization can lead to intragenic transcription, then it would follow that these motifs should be overrepresented in recoded gene regions. To test this hypothesis, we mapped all instances of TANNNT in any frame and compared the number of times they occurred in codon pair deoptimized versus wild-type genes. Recoded *rpoB* contained 3 to 4 times more TANNNT motifs (**Figure 4**). Of note, *rpoB*(2) contained the most TANNNT motifs, which is consistent with the observation that this variant also expressed the greatest number of smaller protein isoforms (**Figure 1B**). We also examined where TANNNTs occurred in relation to the TSS inferred from 5′ end sequencing (**Figure 4**). Several TANNNT motifs in recoded regions appeared prior to one or more iTSS, as expected if our hypothesis were true. For example, only *rpoB*(2) contained two new TANNNT motifs overlapping at nucleotide position 1951 and 1956, corresponding with downstream iTSS at 1963 and 1969. These observations once again support the hypothesis that recoding the codon pair bias of genes enables new iTSS because specific codon pairs give rise to new TSS motifs.

### Mapping protein N-termini confirms translational initiation sites within recoded regions

To explore the relationship between the TSS introduced by recoding and TIS within the resulting transcripts, we detected and mapped the N termini of the additional RpoB protein products. In contrast to studies that map TIS by profiling ribosomal occupancy on a nucleotide sequence, we mapped TIS by detecting the resulting peptide and the results are therefore complementary to ribosome footprinting. Proteins generated from wild-type and recoded *rpoB* variants were enriched by anti-FLAG antibody affinity prior to N-terminal labeling by dimethylation and tryptic peptide detection by mass spectrometry. Several considerations were made in identifying N-termini as bona fide TIS (see Methods for details). For the purposes of this study, we included N-termini that mapped to plausible start codons such as canonical start codons AUG and GUG, or the rarer start codons UUG and CUG (30,31).

We mapped detected TIS to the RpoB sequence. For reasons of data quality, results from *rpoB*(1) were not included in our final analysis, but ample comparisons were still available based on *rpoB*(2) and *rpoB*(3) The resulting maps of N-terminal peptides confirmed that more TIS were detected in recoded gene segments than in the corresponding regions of the wild-type sequence (**Figure 4**). The relative numbers of detected TIS in recoded regions corroborated the multiple isoforms detected by immunoblot (**Figure 1B**). TIS at residues 97, 121, 148, 534, 983, 1045 were detected robustly across the wild type, *rpoB*(2), and *rpoB*(3). However, 30 and 19 TIS were found from the recoded variants *rpoB*(2) and *rpoB*(3) versus 13 TIS in the wild type. Of note, the position of the TIS corresponded qualitatively with recoded regions containing dense TSS (**Figure 4**). Importantly, the higher number of TIS arising from the recoded regions of variant genes was not due to an increase in the absolute number of start codons, since codon usage was largely preserved during recoding. Indeed, the variation in the number of start codons among *rpoB* and recoded variants is low (ranging from 225 to 230 total AUG, GUG, UUC, CUG across the 4 sequences) and thus cannot account for the changes in protein expression.

What do the detected TIS tell us about the start codons that initiate smaller proteins from recoded genes? First, TIS unique to recoded genes appeared more likely to correspond to less common start codons. In wild-type RpoB, only 23% (3 of 13) TIS initiated at the rarer start codons UUG or CUG (compared to the canonical start codons AUG or GUG). In contrast, 57% (12 of 21) and 50% (6 of 12) TIS in *rpoB*(2) and *rpoB*(3) TIS initiated at UUG/CUG. This trend was not an artifact of codon usage changes, as all genes contained similar numbers of in-frame UUG/CUG codons (80, 79, and 73 in the wild type, *rpoB*(2) and *rpoB*(3)) Thus, the frequency of UUG/CUG that functioned as start codons was higher in recoded variants (11% and 8% in *rpoB*(2) and *rpoB*(3)) than in the wild type (4%). From this data, we inferred that recoding likely affected gene expression mechanisms upstream of translation initiation—mechanisms that would boost the accessibility of start codons such as UUG/CUG to an initiating ribosome.

If changing the start codon does not as a rule disrupt or induce translation initiation, what other nucleotide alterations drive changes in TIS? Shine-Dalgarno (SD) sequences are generally thought to be major drivers for translation initiation in bacteria. We hypothesized that synonymous changes that introduce or disrupt intragenic SD-like motifs contributed to the observed TIS. We therefore analyzed the gene sequences for AAG or GGA in the 25 nt preceding each detected start codon. However, we could not discern any obvious relationship between TIS and SD-like sequences. On the one hand, SD motifs were not necessary, as we observed TIS with and without such motifs in both wild-type and recoded segments. On the other hand, both wild-type and recoded genes contained in-frame, intragenic ATG or GTG codons that were preceded by a SD motif but were not detected as a TIS. These results indicate that SD motifs are neither necessary nor sufficient for intragenic translation initiation.

### Transcription initiation motifs are embedded in rare codon pairs

Because gene codon pair score does not correlate with the number of proteins produced, we hypothesized that recoding causes intragenic transcription initiation because particular rare codon pairs encode TSS motifs. To test this hypothesis, we selected codon pairs containing TANNNT, which is the −10 element most frequently observed at endogenous TSS in both *Mtb* and *Msm* (28,29). codon pairs encoding TANNNT were indeed significantly rarer than all other codon pairs (**Figure 5**). Since transcription is a frame-independent phenomenon, the second and third reading frames of a coding sequence should also show selection against enhancer motifs. We therefore calculated the codon pair scores in frames 2 and 3 (strictly speaking, these are triplet pair scores calculated for hexamers, since these frames are generally not translated and thus the term codon does not apply). As predicted, hexamers including TANNNT were also depleted in frames 2 and 3 (**Figure 5**).

**Figure 5.**
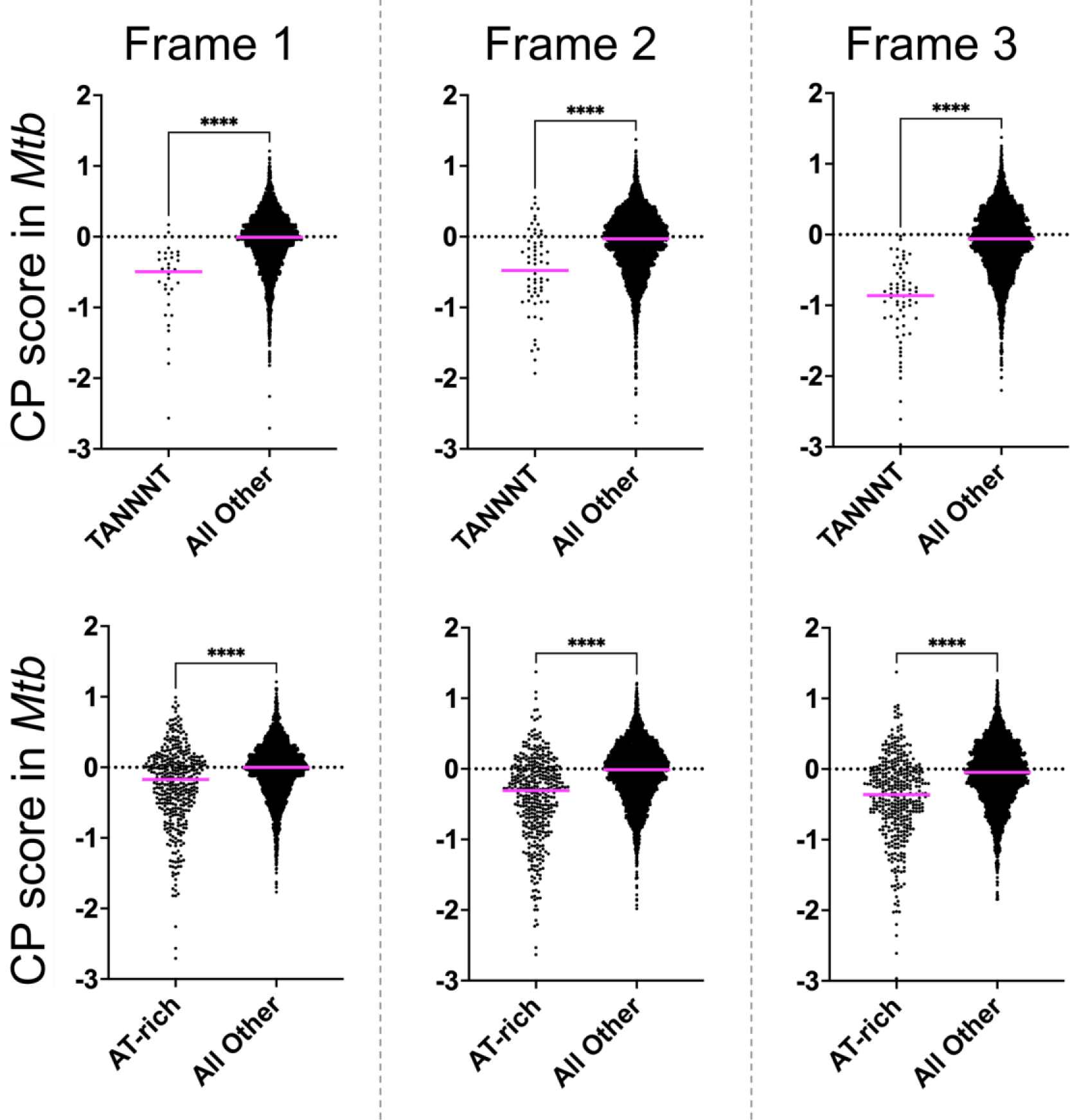
AT-rich codon pairs are depleted in all three frames across the *Mtb* coding genome. Codon pair scores were calculated for each frame (in-frame, frame+1, frame+2) in the *Mtb* coding genome. The scores for codon pairs (A) encoding TANNNT or (B) enriched for A/T (≥4 A/T per codon pair) were compared to all others. Data and average are shown. Unpaired t-test was performed to determine statistical significance (**** p<0.0001).

## DISCUSSION

We set out to attenuate protein expression by increasing the frequency of rare codon pairs in *Mtb* genes by synonymous mutation. Instead, we found that synonymous gene recoding yielded promiscuous intragenic transcription initiation. The resulting mRNAs in turn yielded stable protein products with diverse and highly reproducible sizes and yields. These phenomena were also observed when synonymous recoding was random rather than explicitly designed to introduce more rare codons.

We found that recoding does not consistently attenuate full-length protein expression in mycobacteria (**Figures 1E-G, 2B**). This is contrary to our original hypothesis and to previous publications in codon pair deoptimization that reported the attenuation of fitness in viruses and gene expression in yeast (6,7). The effects of modulating codon pair bias– and making synonymous mutations in general– have not been systematically and broadly examined in bacteria. Overall, the relationship between codon pair bias score and gene expression on the RNA and protein levels needs a detailed examination. Indeed, optimizing codon pair bias has yielded conflicting reports in viruses: Optimization in one case increased, but in another case decreased, protein expression (6,32). The lack of a consistent correlation between gene expression and codon pair bias on the gene (or genome) level is consistent with our conclusion that changes in expression depend instead on the introduction of individual codon pairs.

To our knowledge, this is the first report demonstrating that synonymous gene recoding can give rise to intragenic gene expression in any organism. Our work clearly demonstrates that synonymous mutations can affect not only how proteins are translated but how mRNA is transcribed. This finding undermines the long-standing assumption that synonymous mutations are silent. The idea that synonymous mutations can cause phenotypes is not new (1,33–37). Others have established that synonymous mutations recoding single codon usage can affect mRNA folding and accessibility (36), longevity (38), and rates of translation (39,40). Also, rare codon pairs may attenuate genes in part by causing inefficient translation elongation (6–10). More specifically, some codon pairs cause structural changes in the mRNA that slow down translation by stalling the ribosome at those junctures (8). Other studies have implicated post-transcriptional deficits such as decreased mRNA stability or increased mRNA decay (9,41). Prior research in codon pair deoptimization has been performed predominantly in eukaryotic systems, which might lead to speculation that intragenic protein expression is a mycobacteria-specific phenomenon. However, the dearth of data supporting expression of smaller proteins could plausibly result from conventions in reporting and detection using native antibodies with unknown epitopes (vs. our use of a C-terminal-specific epitope).

Here we found that synonymously recoding genes gave rise to intragenic, functional TSS and posit two explanations for this outcome. First, recoded sequences may boost initiation at endogenous internal transcription start sites (iTSS) that are not otherwise active because they are weaker or only function in specific environmental conditions (*i.e.*, they are cryptic). Indeed, transcriptome-wide TSS mapping studies in mycobacteria have aligned iTSS to two of the genes selected for this study, *rpoB* and *ndh*. *Mtb rpoB* (*Rv0667*) may contain up to 10 different iTSS in addition to 2 TSS in the promoter region that can account for full-length protein (14). The *Mtb ndh* gene is also thought to contain one iTSS, as identified by two different studies (14, 15); none have yet been identified iTSS in *mmpL3*. The prevalence of iTSS in endogenous genes leaves open the possibility that, rather than forming new transcription sites, synonymous recoding is allowing or augmenting transcription initiation at existing cryptic iTSS.

Alternatively, recoding may create entirely new TSS, for example, by introducing new intragenic promoter elements. The particulars of mycobacterial gene expression may make the mycobacterial genome more permissive not only to new TSS, but also transcripts that are translated and therefore functional. Although consensus sequences at both −35 and −10 are generally thought to regulate transcription initiation (42), *Mtb* and *Msm* lack a −35 consensus sequence and may therefore have a lower threshold for initiating transcription (27–29). Also, any new transcripts have a higher likelihood of leading to protein: Mycobacteria readily translate transcripts that begin with just a start codon and up to a quarter of mycobacterial mRNAs are leaderless and lack either a 5′ untranslated region or SD motif (27–29). As we noted above, intragenic or internal TSS (iTSS) are commonly found in mycobacteria, indicating that mechanisms already exist to allow intragenic expression. Indeed, promoters may not be difficult to create in bacteria generally. From a library of randomly generated 100-nt sequences, up to 10% are likely to contain a functional promoter in *E. coli* (43). More recently, Rodriguez et al. showed that synonymous recoding of a marker gene led to a new iTSS and boosted expression from an existing TIS in the antisense direction (44). These observations suggest that promoters readily arise in bacterial genomes and that even less extensive synonymous mutations than the ones we made here are sufficient to create functional promoters.

On the protein level, a surprising result from our data was the detection of numerous N-terminal peptides that would have originated from within the ORF of wild-type *rpoB*. Intragenic TISs in *Mtb* are not unprecedented: Translation initiation is known to occur intragenically in mycobacteria and other bacteria (45). Smith *et al.* inferred by ribosome footprinting that intragenic translation initiation is pervasive in *Mtb* and predicted 4 TISs in endogenous *rpoB* that mapped to RpoB residues 154, 534, 718, and 867 (46). Our results provide direct evidence on the protein level for one of these sites via detection of an N-terminus at residue 534. However, we also detected an additional 12 TIS resulting from wild-type *rpoB* that were not inferred by Smith et al., 10 of which were confirmed by detection from either *rpoB*(2) or *rpoB*(3). The difference in detected TIS in our work vs. Smith *et al.* may indicate the differences in sensitivity between methods used to report TIS and supports the use of multiple approaches to study these sites.

A central finding of our work is that recoding mycobacterial genes causes iTSS. Though our study design did not allow us to catalog all rare codon pairs that can result in iTSS, we confirmed that synonymously recoding genes to contain more rare codon pairs led to aberrant gene expression. Focusing on known transcription initiation motifs, we showed that codon pairs containing TSS motif “TANNNT” or generally TA-rich sequences are rare throughout the *Mtb* coding genome. Further, the TANNNT hexamer is rare in all three frames. Taken together, these data not only offer an explanation for how codon pair deoptimization can cause new ITSS, but also suggest that codon pair biases– and, more broadly, codon context biases– have evolved to curb pervasive transcription in coding regions. We speculate that such biases may serve to help cell machinery discriminate between coding and non-coding sequences.

Ultimately, does the expression of smaller proteins from intragenic start sites have demonstrable biological consequences? While answering this question is beyond the scope of the current study, we note that the isoforms are probably not toxic since we did not observe any obvious changes in bacterial growth for strains containing recoded variants relative to empty vector or wild-type controls. Since all strains used in this study contained the endogenous copy of the recoded gene, we do not know whether the smaller isoforms can modulate the function of the full-length protein. It is tempting to speculate that, independent of physiological function in bacteria, smaller protein isoforms could enhance antigen presentation by the host and provoke an enhanced specific immune response.

Our discovery that synonymous mutations can trigger new transcription may inform best practices in genome annotation, gene engineering, and recombinant genetics. A precise understanding of how nucleotide sequences encode cues for gene expression is critical for accurate genome annotation. Next-generation sequencing technologies continue to rapidly expand genome repositories, but the utility of these data is predicated on the accurate identification of promoters and open reading frames. We provide evidence that bacterial promoters readily arise within open reading frames from synonymous mutations, and this lowers the previously held threshold for transcription initiation (and translation initiation), in agreement with a growing body of research showing that transcription initiation is a relatively promiscuous process in mycobacteria. In addition, codon context optimization has been reported to impact protein production to a greater extent than codon usage optimization alone in both prokaryotes and eukaryotes (47–49). Understanding the relationship between codon pairs and codon context more generally in thus necessary to improve recombinant protein research.

Synonymous mutations are broadly accepted as benign and, by that logic, have been categorically excluded in drug-resistance screens and studies. A recent report on synonymous mutations in drug-resistant *Mtb* showed that, though less common than non-synonymous drug-resistance mutations, a number of synonymous mutations are associated with drug-resistance phenotypes (50). Of note, others have recently demonstrated that synonymous mutations are non-neutral: In a yeast study that characterized 450 mutated variants of 21 genes, synonymously mutating 150 nt-segments tended to confer fitness deficits in 16 of the 21 genes tested (51). Though the exact mechanisms for the non-neutrality of those synonymous mutations in yeast is yet unclear, we have begun addressing here the mechanisms of the unintended consequences for synonymous mutations. Delving further into the mechanisms by which synonymous mutations can disrupt normal gene expression may lend greater insights into how bacteria may use these so-called silent mutations to alter phenotype and perhaps even to develop drug resistance.

This report is the first to offer mechanistic insight into the attenuation of bacterial pathogens by codon pair deoptimization. In doing so, our results inform the implementation of dinucleotide and codon biases to develop live-attenuated vaccines. Especially given the rise of drug-resistant bacteria, vaccine development is an increasingly vital strategy in combating bacterial infectious diseases. Although full-length protein expression was not consistently attenuated, synonymous recoding of essential genes may nevertheless impact mycobacterial survival, pathogenicity, and immunogenicity. Further studies of the biological mechanisms by which codon pair preferences influence bacterial transcription, translation, and physiology will inform the application of this intriguing vaccine strategy.

## MATERIALS AND METHODS

### Bacteria Strains and Culture Conditions

Experiments were performed in *M. smegmatis* (*Msm*) mc^2^155 (ATCC 700084) or *M. tuberculosis* (*Mtb*) mc^2^6020 (H37Rv Δ*lysA* Δ*panCD*; gift of William Jacobs) (**Supplemental Table S1**) (52). *Msm* was grown in Middlebrook 7H9 media (HiMedia) supplemented with 1% (w/v) casamino acid (BD), 0.2% (v/v) glycerol, 0.2% (w/v) glucose, 0.05% (v/v) Tween80 or 7H11 agar (HiMedia) supplemented with 10% (v/v) OADC, 0.5% (v/v) glycerol, 0.05% (v/v) Tween80. *Mtb* was grown in Middlebrook 7H9 supplemented with 10% (v/v) oleic acid/dextrose/catalase supplement (OADC; BD), 0.2% (v/v) glycerol, 24 mg/L pantothenate, 80 mg/L L-lysine, and 0.025% (v/v) Tyloxapol or 7H11 agar supplemented with 10% (v/v) OADC, 0.5% (v/v) glycerol, 0.2% (w/v) casamino acid, 80 mg/L lysine, 24 mg/L pantothenate. NEB5-alpha Competent *E. coli* (C2987, NEB) was used to propagate plasmids and was cultured in LB medium or grown on LB agar. All liquid cultures were grown at 37 °C with shaking at 250 rpm for *Msm* and 110 rpm for *Mtb*. Strains containing selectable markers were cultured on medium with 50 µg/mL hygromycin (Hygromycin B Solution, Mirus) or 25 µg/mL kanamycin (GoldBio) for mycobacterial strains and 100 µg/mL hygromycin or 50 µg/mL kanamycin for *E. coli*. See **Supplementary Table S2** for a list of key reagents.

### Calculation of codon pair scores

Complete H37Rv genome collected (per Mycobrowser 3/23/2021 release, acquired 6/30/2021) for processing; non-coding and RNA-encoding genes were excluded from analysis (53). First, the number of times any codon pair occurred in-frame in all coding sequences were recorded as the observed frequency of each codon pair. Next, the expected frequency was determined using an algorithm that controlled for codon usage as well as the distribution of rare and common codons across coding genes. The algorithm randomly shuffled synonymous codons within each gene to generate 10,000 synonymous random-codon-shuffled sequences. The number of times a codon pair was observed in a random-shuffle sequence was then averaged across all 5000 iterations (sum of all codon pair frequencies / number of iterations). The number of occurrences of a codon pair in a given random-codon-shuffle was then totaled across the entire coding genome and averaged by the total number of iterations (*i.e.*, 10,000). This value (the average number of times a codon pair occurs across 5000 randomly shuffled sequences) is an approximation for the expected frequency of a given codon. A Codon Pair Score (codon pairS) was calculated for each hexameric codon pair by the following formula: 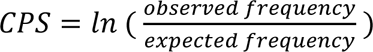.

### Generating Recoded Variants

Codon pair deoptimized genes were designed by synonymously recoding a length of *Mtb rpoB* (gene locus Rv0667), *mmpL3* (Rv0206c), or *ndh* (Rv1854c) to maximize the number of deoptimized codon pairs. Recoded regions were codon-pair deoptimized in a similar manner to previously published methods (54). Briefly, codon pairs were shuffled synonymously at random one million times within each recoded segment of *rpoB*, *mmpL3*, or *ndh*. Shuffles that improved the number of rare codon pairs were selected in order to generate deoptimized sequences. For the *rpoB* deoptimization score variants (range of codon pairB scores) used in the score variation study, deoptimized sequences were selected that matched a designated range of codon pairB scores (codon pairB = 0, 0.01, 0.02, or −0.05). For the maximally deoptimized experimental variants (*rpoB*(1), (2), (3); *mmpL3*(1), (2), (3); *ndh*(1), (2)), recoded codons were reverted to the wild type at positions where the original codon offered a comparable or higher codon pair score.

### Molecular Cloning

For a list of all vectors and primer sequences, see **Supplemental Tables S3, S4**. Wild-type genes *rpoB*, *ndh*, and *mmpL3* were amplified from *Mtb* H37Rv genomic DNA by PCR and inserted into pGW1-6C (gift of Tom Alber, University of California, Berkeley) by InFusion (Takara Bio) for wild-type genes or by ligation for recoded gene variants using restriction sites SpeI and NdhI. Recoded gene variants were obtained by synthesis and cloned into pUC57 by Genscript. Wild-type and recoded genes were also subcloned from pGW1-6C into the integrating vector pMV306 and episomal pMV261 (25) via the XbaI and ClaI restriction sites using ligation or InFusion. All restriction enzymes and T4 ligase were purchased from New England Biosciences.

The *rpoB* codon pair score variants with codon pairBs of 0, 0.01, 0.02, or −0.05 were synthesized in ∼1.2 kb fragments (Twist Bioscience) and assembled by PCR into a final product of ∼3.6 kb each. Variants were subsequently cloned into pGW1-6C by Gibson Assembly (NEB). Nonsense mutations were generated by site-directed mutagenesis using primers onk046 and onk047 to change the *rpoB* start codon TTG to stop codon TAG. Deletion mutants for *mmpL3* and *ndh* were similarly cloned using site-directed mutagenesis primers designed to delete the first 60 nucleotides of the open reading frame. Transcription terminator constructs were cloned from the pGW1-6C constructs by restriction digest with XbaI and SpeI to first remove the promoter. The plasmid was re-circularized using primers omlp759 and omlp760 to insert the *E. coli rrnB* T1 terminator or omlp761 and omlp762 to insert a T2 terminator (26).

### Generation of Bacterial Lysates

*M. smegmatis* was grown in 10-mL cultures for immunoblotting or 100-mL cultures for affinity purification. Cells were pelleted by centrifugation at 3,000 *g* for 10 min and washed in sterile PBS or TBS as noted further below. Decanted pellets were stored at −80 °C and thawed just prior to lysis under buffer conditions described in further detail in following sections. After thawing on ice, cells were resuspended in buffer and lysed with 0.1 mm zirconia beads (BioSpec) by bead beating (Beadruptor, Omni International) for 30 s at 6 m/s followed by 5 min incubations on ice for a total of 3 cycles. Lysates were cleared by centrifugation at 5,000 x*g* and protein concentration was quantified by Pierce BCA Protein Assay (Thermo Fisher).

### Immunoblotting

Thawed and washed cell pellets were lysed in 1 mL sterile PBS with 10% (v/v) glycerol and 1x cOmplete mini protease inhibitor cocktail (Roche). Ten micrograms of protein were separated by SDS-PAGE and transferred to a nitrocellulose membrane (Bio-Rad). Membranes were probed with α-FLAG M2 (Sigma-Aldrich F1804, 1:1,000 dilution) or α-GroEL (Santa Cruz #5177, 1:5,000), followed by IRDye 800CW Goat anti-Mouse IgG secondary antibody (LI-COR, 1:15,000) and imaged (Odyssey Cx scanner, LI-COR).

### RNA Isolation

*M. smegmatis* was grown in 10 mL cultures overnight to optical density at 600 nm (OD_600_) of 1-1.5. Cells were pelleted by centrifugation, resuspended in 1 mL of TRIzol LS (Invitrogen), and stored at −20 °C. TRIzol resuspensions were thawed on ice before lysis by bead beating (Beadruptor, Omni International) for 30 s at 6 m/s followed by 5 min on ice for a total of 3 cycles. Total RNA was extracted from the aqueous phase by isopropanol precipitation. RNA prepared for qPCR was then purified using columns according to the manufacturer’s instructions (Qiagen RNeasy kit). Precipitated RNA for 5’ RACE or column purified RNA for qPCR was then treated with DNase using a commercial kit (Turbo DNAse, Ambion, AM2238). Total RNA yield was verified on a 2% agarose gel by visual inspection. To confirm the removal of DNA, DNase-treated samples were used as templates for qPCR targeting 16S rRNA using primers onj005 and onj006, and sample purity was checked by comparing absorbance at 260 vs. 280 nm (ND-1000, Thermo Fisher).

### qRT-PCR Analysis of Terminator-Driven *rpoB* Variants

RNA samples were used as template for cDNA synthesis with random primers (Verso Kit, Thermo Scientific). The cDNA libraries were diluted 10-fold and 2 µL of the dilution was used for qRT-PCR analysis (Power SYBR Green PCR Master Mix, Applied Biosystems) on a LightCycler 480 Instrument (Roche). Ct values were calculated by LightCycler 480 Software (Roche) and normalized using 16S rRNA as a housekeeping gene. Primer efficiencies were verified by standard curves. Gene expression fold-change was calculated using the delta-delta Ct method (Livak and Schmittgen, 2001).

### Construction of 5’-End-Mapping Libraries

5’-End-Mapping libraries were constructed using a protocol similar to those previously published by others (29,59,60). RNA was isolated from *Msm* expressing 5’ transcription terminator *rpoB* as described above. DNase-treated RNA was ligated with a 5’-hydroxylated ‘Processed Start Site’ (PSS) RNA adaptor oligomers (onkh031) by RNA Ligase 1 (Promega or NEB) and then purified to remove excess oligomers (NEB Monarch RNA Cleanup kit). RNA was then treated with RNA pyrophosphatase (NEB) and then ligated with a ‘Transcription Start Site’ (TSS) RNA adaptor (onkh032). PSS- and TSS-labeled RNA was then used as a template for cDNA synthesis by random primers (Verso Kit, Thermo Scientific).

For 5’ mapping, 5’-ends were amplified by tiling PCR using a TSS adaptor-specific forward primer (onkh037) and one of 7 gene-specific reverse primers with annealing sites spanning the length of *Mtb rpoB* wildtype or variants at ∼500 bp intervals (**Supplemental Table S4**). A total of 7 PCR reactions were thus performed using each cDNA library template, resolved on a 1.8% agarose gel. Single bands were cut and gel purified for Sanger sequencing (Genewiz) for preliminary validation.

After confirming several intragenic TSS by Sanger sequencing, 5′ RACE PCR products were repeated, this time running each *rpoB*-specific reverse primer in two reactions with the TSS tag-specific forward primer or the PSS tag-specific forward primer onkh175 for a total of 14 reactions per template (one of 2 forward primers onkh037 or onkh175 with each of 7 reverse primers). PCR products were pooled by template and purified (Nucleospin, Takara). RACE amplicons were end repaired, A-tailed, and an Illumina compatible barcode was ligated using the Roche Kapa HyperPlus DNA Library Preparation kit. Individually barcoded libraries were amplified with Kapa HiFi HotStart DNA Polymerase and quantified with Invitrogen Qubit dsDNA High Sensitivity Assay. Library quality was validated with Agilent Bioanalyzer to determine the appropriate bp size. Libraries were pooled and quantified by qPCR with Kapa Illumina/Universal Library Quantification Kit. The pooled library was then sequenced on an Illumina MiSeq (PE150 with index read).

### Sequencing Analysis

Sequence files (FASTQ) were trimmed for Illumina universal adapters with Cutadapt v4.0 (61). Quality control was done using FastQC v0.11.8 (62). Reads were selected for 16 bases start sites (TSS or PSS) in the beginning of read 1 with 2 mismatches allowed using SeqKit v2.2.0 (63). We removed the start site sequences from the first 16 bases and aligned trimmed sequences to the rpoB gene (wild-type and 3 variants) using STAR v2.7.8a with default parameters (64). Unique alignments were selected by SAMtools v1.14 (65). Downstream statistical analysis was performed using basic functions and modules from Python v3.8.8, pandas v1.2.4 (66,67), and Microsoft Excel.

We thus compiled all sequences preceded by a start tag (TSS or PSS) that also aligned to one of our genes of interest as follows (68): First, we normalized the TSS hit list against the list of PSS-tagged 5’-ends, retaining only the ends that had were 10-fold or more abundantly TSS-tagged *vs.* PSS-tagged. 5’-ends tagged with predominantly PSS sequences were eliminated from our hit list as processed starts. Of the remaining hits, we further selected only 5’-ends with the highest coverage in a 10 nt window to eliminate any redundant start sites resulting from imprecise transcription initiation (68). This was done prior to inferring transcription start site vs. processed sites from the 5’-end of an aligned sequences (**Supplemental Tables S5-S7**).

### Protein Affinity Enrichment

Thawed and washed cell pellets were lysed in 3 mL sterile TBS with 10% glycerol, 1% Triton X-100, protease inhibitor cocktail (Roche). After protein quantification, whole cell lysates were precleared as follows: Lysates were incubated on a rotator at room temperature for 1 hour with 7 µg/mL final concentration Immunoglobulin G1 (Sigma-Aldrich M5284) and then affinity purified with Protein G Magnetic Beads (Pierce). Briefly, beads were equilibrated according to the manufacturer’s instructions. 1/100^th^ lysate volume of resin was added to the lysate/IgG mixture and incubated at room temperature for 1 h with gentle mixing. After the beads were collected with a magnetic separator, the supernatant was removed and used as precleared input for affinity purification with 1/30^th^ volume α-FLAG M2 Magnetic Beads (Millipore M8823). Beads and supernatant were incubated on a rotator overnight at 4°C. The resulting supernatant was collected and the beads were washed three times with 20X resin volume TBS. Elution buffer was prepared immediately before use by adding 3XFLAG peptide (Sigma F3290) to TBS to a final concentration of 150 ng/µL. Proteins were eluted by incubating with 2X resin elution buffer for 15 min at 22 °C with rotation. This elution step was repeated and eluates were pooled. The final output was validated silver stain (Pierce Silver Stain Kit, Thermo Fisher) and visual inspection prior to processing for mass spectrometry.

Eluates were reduced, alkylated, separated on SDS-PAGE gel. All protein bands in each sample were pooled by excising the entire gel lane, followed by N-terminal labeling with triethylammonium bicarbonate and trypsin in-gel digestion. An aliquot of the resulting desalted peptides was analyzed with an EASY-nLC 1200 coupled to a Thermo Fisher Q Exactive HF-X Mass Spectrometer operated in data dependent mode. The resulting spectral data were searched against *M. tuberculosis* RpoB wild-type and recoded sequences using Byonic and N-terminally modified peptides were verified manually.

### Inferring Start Codons from N-terminal Labeled MS/MS Hits

We compiled all spectra showing an N-terminal diMe, Ac, or fMet. We excluded any spectra falling below a threshold Byonic score of 300, which measures the quality of a given peptide-spectrum match (55). Any diMe or Ac residues immediately following an R or K trypic cut site were excluded to avoid false positives. All hits indicating diMe-M or Ac-M were also excluded given that any N-terminal Met would be expected to be fMet. All hits indicating diMe-K were excluded because of possible side-chain modification. In processing the Ac hits, all hits that followed a K or R residue were excluded. Despite evidence that Ac-T and Ac-S are frequently found at the N-terminus, we also excluded those amino acids because of possible side-chain modifications, as well as Ac-K, Ac-Y, and Ac-H (56–58). Finally, N-terminal residues were inferred to represent a true translation initiation event if the peptide position mapped to a codon +1 from a start codon (**Supplemental Table S8**). For the purposes of this study, we accepted ATG, GTG, TTG as typical start codons, as well as the less common start CTG (30,31).

## ACKNOLWEDGMENTS

Our deepest thanks to Todd Gray for experimental advice and thoughtful discussions. Thanks also to the Seeliger lab for helpful discussions. This work was supported by R35 GM128552 (J.C.S.). N.K.H. was also supported by T32 GM008444.

## SUPPLEMENTAL INFORMATION

**Supplemental Figure S1.** The CP bias of a coding sequence is scored as the mean of each CP.

**Supplemental Figure S2.** Expression of smaller proteins from recoded genes are not an artifact of gene copy number or selection with the translational inhibitor hygromycin

**Supplemental Figure S3.** Substituting a TAG stop codon for the start codon does not abolish expression of full-length Ndh or MmpL3

**Supplemental Table S1-S4.** (S1) Strains, (S2) Reagents, (S3) Plasmids and (S4) Primers used in this study.

**Supplemental Table S5.** Analysis of TSS for wild-type *rpoB*

**Supplemental Table S6.** Analysis of TSS for *rpoB*(2)

**Supplemental Table S7.** Analysis of TSS for *rpoB*(3)

**Supplemental Table S8.** Analysis of TIS for *rpoB* wild type and variants (1), (2), and (3)

