## Supplemental Figures S1-S3 for "Gene recoding by synonymous mutations creates promiscuous intragenic transcription initiation in mycobacteria"

**Supplemental Information**

- **Figure S1.** The CP bias of a coding sequence is scored as the mean of each CP.
- **Figure S2.** Expression of smaller proteins from recoded genes are not an artifact of gene copy number or selection with the translational inhibitor hygromycin
- **Figure S3.** Substituting a TAG stop codon for the start codon does not abolish expression of full-length Ndh or MmpL3

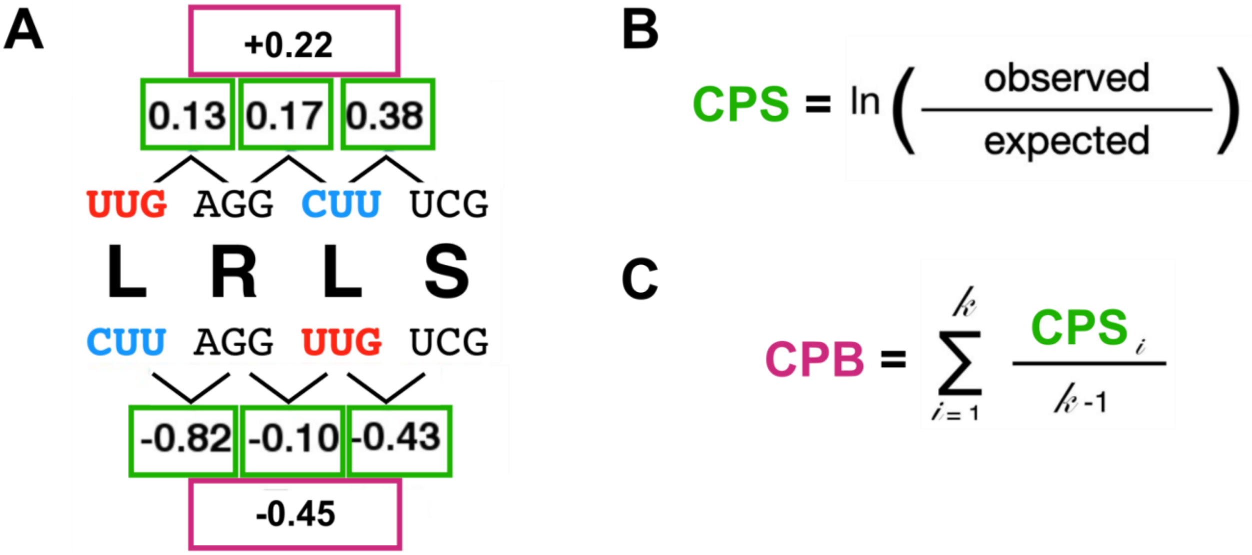


**Supplemental Figure S1. The CP bias of a coding sequence is scored as the mean of each CP.** A short coding sequence encoding Leu-Arg-Leu-Ser can be coded two ways, shown above and below the peptide sequence in panel (A). Each Leu is encoded by a unique, synonymous codon, UUG or CUU. The position of these two Leu codons are switched in the top and bottom nucleotide sequences, effectively changing the CP biases even though the codon usage is the same. (B) The CP score (CPS, in green text) is determined by calculating the natural log of the ratio of the total number of times a given CP is observed in an organism's coding genome to the number of times the CP is expected to appear, controlling for codon usage (refer to methods). Examples of CP scores in Mtb are boxed in green above the corresponding CP in panel (A). (C) The CP bias score (CPB, text in purple) assigned to a given gene is the arithmetic mean of all the CPS found in the sequence. The mean of the CPS in panel (A) have been thus averaged, and the final CPB of the top and bottom nucleotides appear boxed in purple. Figure adapted from Coleman et al. 2008.

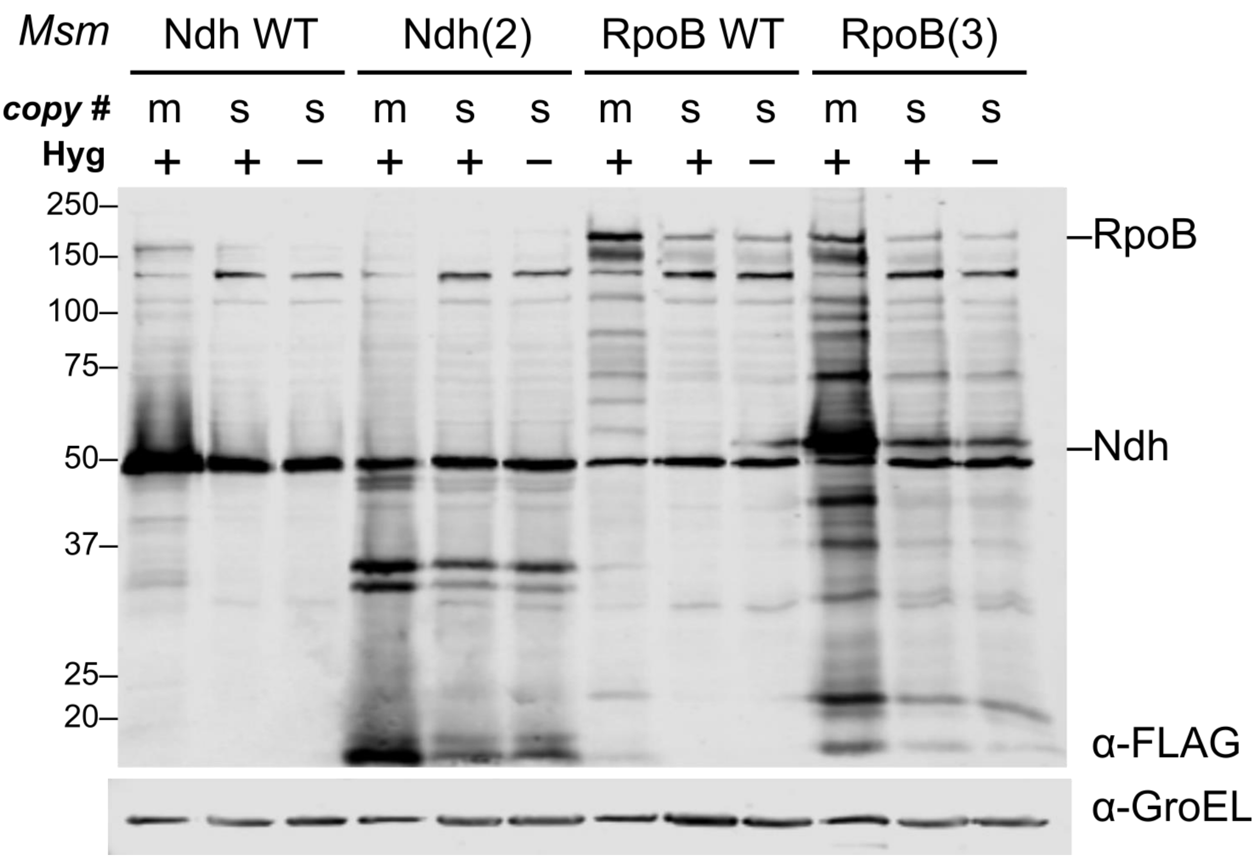


**Supplemental Figure S2. Expression of smaller proteins from recoded genes are not an artifact of gene copy number or selection with the translational inhibitor hygromycin.** Immunoblot of wild-type and recoded Mtb genes expressed in Msm multicopy episomal ("m") or single copy integrated ("s") plasmids. Strains with stably integrated plasmids ("s") were grown with or without hygromycin, as indicated. Blot is representative of 3 biological replicates. GroEL blot is included as a loading control.

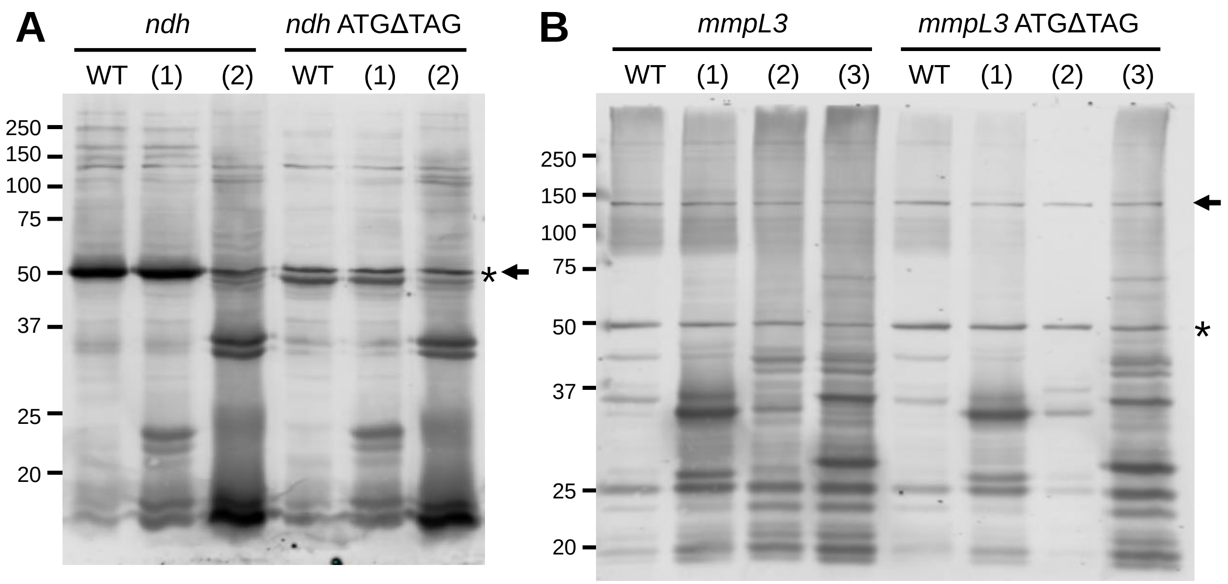


**Supplemental Figure S3. Replacing the start codon with the TAG stop codon does not abolish expression of full-length Ndh or MmpL3.** Anti-FLAG immunoblots of total lysates from *Msm* expressing the designated wild-type or recoded (A) *ndh* or (B) *mmpL3* constructs without (*left*) or with (*right*) the start codon substituted with the TAG stop codon (ATGΔTAG). Arrows designate the expected migration of full-length protein. Asterisks denotes a non-specific FLAG cross-reacting band in *Msm* total lysates. This band is nearly coincident with full-length Ndh (see also Figure 1I). Immunoblots are representative of 3 (*ndh*) or 2 (*mmpL3*) independent experiments.
